# Reduction of PALS1/Nok disrupts retinal lamination through altered cell positioning while preserving photoreceptor self-organization capacity

**DOI:** 10.64898/2026.06.16.732437

**Authors:** Gonzalo Aparicio, Flavio R. Zolessi

**Author notes:** **Corresponding author:** F. R. Zolessi. **Emails**: G. Aparicio.

## Abstract

The vertebrate neural retina is composed of several neuronal types that precisely organize into layers, with photoreceptors facing the outer surface and the projection neurons, retinal ganglion cells, at the innermost layer. This organization, essential for its function, is established during early development through a complex process involving cell-cell interactions such as adhesion. In the case of photoreceptors, two adhesion complexes, based on the adhesive proteins Cadherin2 and Crumbs, appear essential for their correct localization at the outer nuclear layer (ONL). We here aimed at better characterizing the role of the scaffolding protein PALS1, a central component of the Crumbs complex. Through a validated *pals1a*/*nok* morpholino knockdown strategy in zebrafish embryos, we demonstrate that its reduced expression causes photoreceptor progenitors to initially disperse as actively migrating cells, to then coalesce into cell groups around the central retina. They eventually start polarizing, forming rosette-like structures with the apical border towards the inside. Conversely, in organoids derived from uncommitted neuroepithelial retinal progenitors, PALS1 deficiency causes an inversion of their localization from internal rosette-like structures to an organized superficial layer. In both conditions, photoreceptors show signs of polarization, with apical borders towards the inside of rosettes in wild-type organoids, to surface-directed apical borders in morphants. Altogether, our results support previous observations of the pivotal function of the Crumbs complex in ONL formation, but also indicate that either the Crumbs complex, or PALS1 itself, are central for the delicate balance in differential cell adhesion partly responsible for retinal lamination.

## Introduction

Laminar organization is a key feature of the central nervous system in vertebrates, by means of which cell bodies from neurons with different functions are segregated into layers, separated by areas of synaptic contact or axonal extension. The neural retina is one of the clearest examples of this type of tissue organization, where sensory neurons, the photoreceptors, are all restricted to the outer-most layer (the outer nuclear layer, ONL), followed by an inner nuclear layer (INL) with different types of interneurons and glial cell bodies, and a ganglion cell layer where the projection neurons are located (the retinal ganglion cells, RGCs). These structural features are acquired during early development through the gradual differentiation of each cell type, with the RGCs differentiating first, and the ONL separating at the end of the process, as has been studied in detail in the zebrafish (Hoon et al., 2014; Norden, 2023). Since the retina derives from a neuroepithelium, it is not surprising that there is a close relationship between the polarity of this early epithelium and that of the mature organ. In fact, the outer border of the retina is lined by an apical surface, from which photoreceptor inner and outer segments extend. Contrary to other neuronal types, cones begin their morphological differentiation before their last cell division. In fact, *in vivo* imaging of photoreceptor progenitors in the zebrafish embryo has shown how basal process detachment, apical nuclear migration and dense cell packing are events that take place well before the formation of the outer plexiform layer (Aparicio et al., 2021; Rocha-Martins et al., 2023; Suzuki et al., 2013; Weber et al., 2014).

Which cell polarity signals are responsible for this accurate process? Early mutational screens in the zebrafish uncovered the role of several apical adhesion complex proteins in the organization of retinal layers, all mutants having phenotypes characterized by the redistribution of neurons to ectopic locations (Malicki et al., 1996). Most of these mutations were mapped to genes related to epithelial cell polarity proteins, such as components of adherens junction complexes, like *parachute*/*glass onion*/*labyrinth* (*cdh2*, encoding Cadherin2/N-cadherin; Erdmann et al., 2003; Malicki et al., 2003; Masai et al., 2003), the Crumbs complex, like *oko meduzy* (*ome*, encoding Crb2a; Malicki and Driever, 1999) or *nagie oko* (*nok*, encoding PALS1/Nok/MPP5a; Wei and Malicki, 2002), and the Par6 complex, like *heart and soul* (*has*, encoding the atypical PKCζ; Pujic and Malicki, 2004).

The Crumbs complex is highly conserved in metazoan evolution, being usually localized at the apical-most side of epithelial cells, and in continuous crosstalk with the other apical complexes and the Hippo pathway (Buckley and St Johnston, 2022; Martin et al., 2021). Crumbs is a transmembrane protein, with several isoforms encoded by different genes, and has been found to be an adhesion molecule with essential roles in retinal development (Bulgakova and Knust, 2009; Omori and Malicki, 2006). In addition, mutations in *crb* genes in humans are causative of retinal dystrophies, such as retinitis pigmentosa and Leber congenital amaurosis, underlying the biomedical importance of these proteins in retinal integrity (Boon et al., 2020; Quinn et al., 2017). PALS1 is an essential intracellular component of the Crumbs complex and a member of the membrane-palmitoylated protein (MPP) subfamily of membrane-associated guanylate kinases (MAGUK), whose multidomain scaffolds organize apical and lateral protein complexes across a wide range of cell types (Chytła et al., 2020; Roh et al., 2002). There are two ortholog genes encoding PALS1 proteins in the zebrafish, the aforementioned *pals1a*/*nok*/*mpp5a* and *pals1b*/*ponli*/*mpp5b*. The protein product of *pals1a*/*nok*, which we will call from here PALS1, is ubiquitous and co-localizes with the three major Crumbs proteins in the early central nervous system of the zebrafish (Zou et al., 2013). In the differentiating retina, it becomes restricted to photoreceptor progenitors, localizing close to the retinal outer limiting membrane (OLM; Wei et al., 2006; Zou et al., 2013). The protein encoded by *pals1b*/*ponli*, on the other hand, is only expressed in a restricted way, in postmitotic photoreceptors (Zou et al., 2010). Previous work showed that the null mutation of the *pals1a*/*nok* gene causes a very severe retinal phenotype, including disruption of the RPE leading to the contact of the apical side of the neuroepithelium with the RPE basal lamina, the Bruch’s membrane (Wei and Malicki, 2002; Zolessi et al., 2006).

Using *in vivo* confocal imaging of early zebrafish embryos, we previously demonstrated that in *cdh2* knockdown conditions, photoreceptor progenitors transitioned from a dispersed distribution to a rosette-like organization. We demonstrated a dynamic behavior of these photoreceptor progenitors comparable to that of wild-type embryos when they coalesce and polarize at the ONL (Aparicio et al., 2021). In the present work, we aimed at better characterizing the importance of *pals1a*/*nok* regarding photoreceptor early differentiation and the formation of the ONL in the zebrafish, by means of a controlled knockdown approach. By using a combination of live imaging and the generation of organoids from early retinal neuroepithelial cells, we observed that these cells retained the capacity for self-organization in the PALS1-deficient conditions, organizing in rosettes in intact retinas and re-organizing from an internal to a superficial position in organoids. Altogether, our results show an essential role for PALS1 in the regulation of photoreceptor positioning and ONL formation in the zebrafish retina, independently of the mechanisms ensuring cell-type laminar segregation and cell polarization.

## Experimental procedures

### Zebrafish husbandry and transgenic lines

Zebrafish (*Danio rerio*) were maintained and bred using standard methods with a 14/10 h light/dark cycle. Embryos were raised at 28.5° C and staged in hours post-fertilization (hpf) as described (Kimmel et al., 1995). To prevent pigmentation, embryos were incubated from 8 hpf with 0.003 % 1-phenyl-2-thiourea (PTU; Sigma-Aldrich). We used wild-type (Tab-5) and previously established transgenic lines: Tg(*crx*:EGFP-CAAX,*myl7*:EGFP) (Aparicio et al., 2021), Tg(*atoh7*:Gap43-mRFP)^cu2^ (Zolessi et al., 2006), and Tg(*gfap*:EGFP)^mi2001^ (Bernardos and Raymond, 2006). All manipulations were carried out following the approved local regulations (CEUA-IPMon and CNEA, Uruguay).

### Gene knockdown and CRISPR/Cas9 genome editing

Loss of function of *pals1a/nok* was achieved by two complementary approaches: translation-blocking morpholino oligomers (MOs) and CRISPR/Cas9-mediated genome editing in F0. Translation-blocking MOs (Gene Tools, Philomath, OR) were microinjected into the yolk of 1- to 4-cell stage embryos. The *pals1a/nok* MO (Zolessi et al., 2006) or a standard control MO (same concentration as *pals1a/nok* MO) were co-injected with a *p53* MO (1.5 ng) to suppress non-specific apoptosis (Robu et al., 2007); these two conditions are hereafter referred to as *pals1a/nok* MO and Standard MO, respectively. All MO sequences are listed in Supplementary Table 1.

CRISPR/Cas9 F0 mutagenesis was performed following the protocol described in Wu et al., 2018, with four gRNAs designed against the *pals1a/nok* genomic sequence (guide sequences listed in Supplementary Table 2). Briefly, four gRNAs targeting contiguous regions across exons 3, 4 and 6 of *pals1a/nok* were designed using CRISPRscan (Moreno-Mateos et al., 2015). Double-stranded DNA templates were generated by PCR following a cloning-free strategy (Varshney et al., 2015), with a target-specific forward primer carrying the T7 promoter and a generic reverse primer providing the tracrRNA scaffold, and transcribed *in vitro* overnight at 37 °C with the MAXIscript T7 kit (Thermo Fisher Scientific, AM1320). The four gRNAs were pooled and co-injected with HiFi Cas9 Nuclease V3 (IDT, #1081061) as a ribonucleoprotein complex into the yolk of 1- to 4-cell stage embryos (see Supplementary Table 2 for guide sequences and injection details). The phenotypic effects of CRISPR/Cas9 mutagenesis were assessed by comparing F0 crispants with *pals1a/nok* morphants in terms of overall embryonic morphology and retinal organization. Editing at the targeted loci was further confirmed by PCR amplification followed by agarose gel electrophoresis, which revealed the heteroduplex banding pattern indicative of efficient mutagenesis in injected embryos.

### Time-lapse confocal microscopy

Embryos at 36 hpf were dechorionated, anesthetized in 0.04 mg/mL tricaine methanesulfonate (MS-222; Sigma-Aldrich) diluted in E3 medium, and mounted laterally in 1% low-melting-point agarose (Invitrogen, 16520) supplemented with 0.04 mg/mL MS-222 and 0.003% PTU to prevent melanization, in glass-bottom dishes (MatTek). After agarose gelation, dishes were covered with E3 medium containing 0.04 mg/mL MS-222 and 0.003 % PTU. Time-lapse imaging was performed at 30° C using a Zeiss LSM 880 confocal microscope with a 25×/0.8 NA glycerol immersion objective. Z-stacks spanning ∼50 μm were acquired at 1 μm intervals in bidirectional scanning mode with 512×512-pixel resolution every 10 min for 12-15 h, with acquisition time of approximately 1 min per embryo. Up to 8 embryos were imaged per experiment. Laser intensity was maintained at minimum levels required for signal detection.

### Retinal organoid culture

Retinas were dissected from 24 hpf embryos and dissociated following the protocol described by Eldred et al. (2017) with minor modifications. Briefly, dissected retinas were incubated in 0.25 % trypsin-EDTA (Sigma-Aldrich) for 10 min at room temperature, then mechanically dissociated by gentle pipetting in calcium-free medium (116.6 mM NaCl, 0.67 mM KCl, 4.62 mM Tris, 0.4 mM EDTA, pH 7.8, supplemented with 100 μg/mL heparin, 0.003 % PTU, and 1 % penicillin-streptomycin). Agarose microwell dishes were prepared using 35-well PDMS molds (Microtissues). Molds were cast with 2 % UltraPure LPM agarose (Invitrogen) and equilibrated with L-15 medium supplemented with 1 % penicillin-streptomycin. Single-cell suspensions were seeded in a dropwise manner and allowed to settle for 15-20 min before adding culture medium consisting of L-15 medium supplemented with 10 % embryo extract, 3 % fetal bovine serum, 2 % N2 supplement, 0.003 % PTU, and 1 % penicillin-streptomycin. Cell aggregates were cultured at 28° C for 48 h before fixation and analysis. Embryo extract was prepared from dechorionated 72 hpf embryos treated with 0.5 % sodium hypochlorite for 2 min, rinsed in Ringer’s solution for 2 min, and homogenized on ice using a Dounce homogenizer. L-15 medium with 1 % penicillin-streptomycin was added to prepare 1 mL extract per 200 embryos.

### Immunofluorescence

Embryos were fixed overnight at 4° C in 4 % paraformaldehyde in PBS, permeabilized, and subjected to whole-mount immunostaining as previously described (Aparicio et al., 2021). Primary antibodies: anti-GFP (DSHB, DSHB-GFP-12A6, 1:200), anti-Zpr1 (ZIRC, 1:100), anti-zn-8 (ZIRC, 1:200), anti-aPKCζ (SCBT sc-216, 1:500). Secondary antibodies from ThermoFisher Scientific: anti-mouse IgG-Alexa 488 (A11034), 1:1000; anti-mouse IgG-Alexa 633 (A21050), 1:1000; anti-rabbit IgG-Alexa 633 (A21070), 1:1000. F-actin was visualized with TRITC-phalloidin (Sigma-Aldrich, P1951, 0.4 μg/ml). Nuclei were labeled with methyl green (2 μg/mL; Prieto et al., 2014). Samples were mounted in 75 % glycerol in 50 mM Tris buffer, pH 8.0, and imaged on Zeiss LSM 800 or LSM 880 confocal microscopes.

### Image analysis and statistics

Images were processed using Fiji software (Schindelin et al., 2012). Time-lapse experiments were corrected for embryo movement during acquisition using the Correct 3D Drift plugin (Parslow et al., 2014). Cell tracking and quantitative analysis of cell migration were performed using the Manual Tracking and Chemotaxis and Migration Tool plugins. Cell height (apico-basal axis length) of *crx*:GFP-positive cells was measured manually in Fiji on single confocal planes as the distance between the apical and basal ends of each cell. The apico-basal position of *crx*:GFP-positive nuclei was expressed as the distance to the apical surface normalized to total retinal width. Within organoids, photoreceptor-progenitor distribution was quantified as the relative position of each cell along the center-to-border axis (0 = center, 1 = border). Briefly, the organoids 3D centroid was established manually looking through the different confocal stacks. The relative position of the cells was determined by measuring their distance to the center, and dividing it by the radius in each case. Statistical analyses were performed using GraphPad Prism 8. Data normality was assessed using D’Agostino-Pearson or Shapiro-Wilk tests. Comparisons were made using Student’s *t*-test for normally distributed data or Mann-Whitney test for non-normally distributed data. Results are presented as mean ± SD or median with interquartile range as appropriate. Statistical significance was set at p < 0.05.

## Results

### Photoreceptor positioning disruption upon *pals1a/nok* knockdown using MOs and CRISPR/Cas9

With the aim of setting up an experimental situation in which we could test the role of *pals1a/nok* in photoreceptor early differentiation and localization in the neural retina, we decided to test different knockdown conditions using an already characterized MO (Zolessi et al., 2006) and CRISPR/Cas9 in F0, using four custom-designed gRNAs (see Supplementary Table 2), in Tg(*crx*:EGFP-CAAX,*myl7*:EGFP) embryos, from here referred to as Tg(*crx*:GFP). Crispant embryos at 72 hpf showed a wide distribution of *crx*-positive cells (photoreceptor progenitors and differentiating photoreceptors) across the retina (Fig. 1A). Most cells were found in isolation, with others forming large elongated groups eventually subdivided into rosettes, in a phenotype that appeared slightly milder than that described in null mutants (Wei et al., 2006).

**Figure 1.**
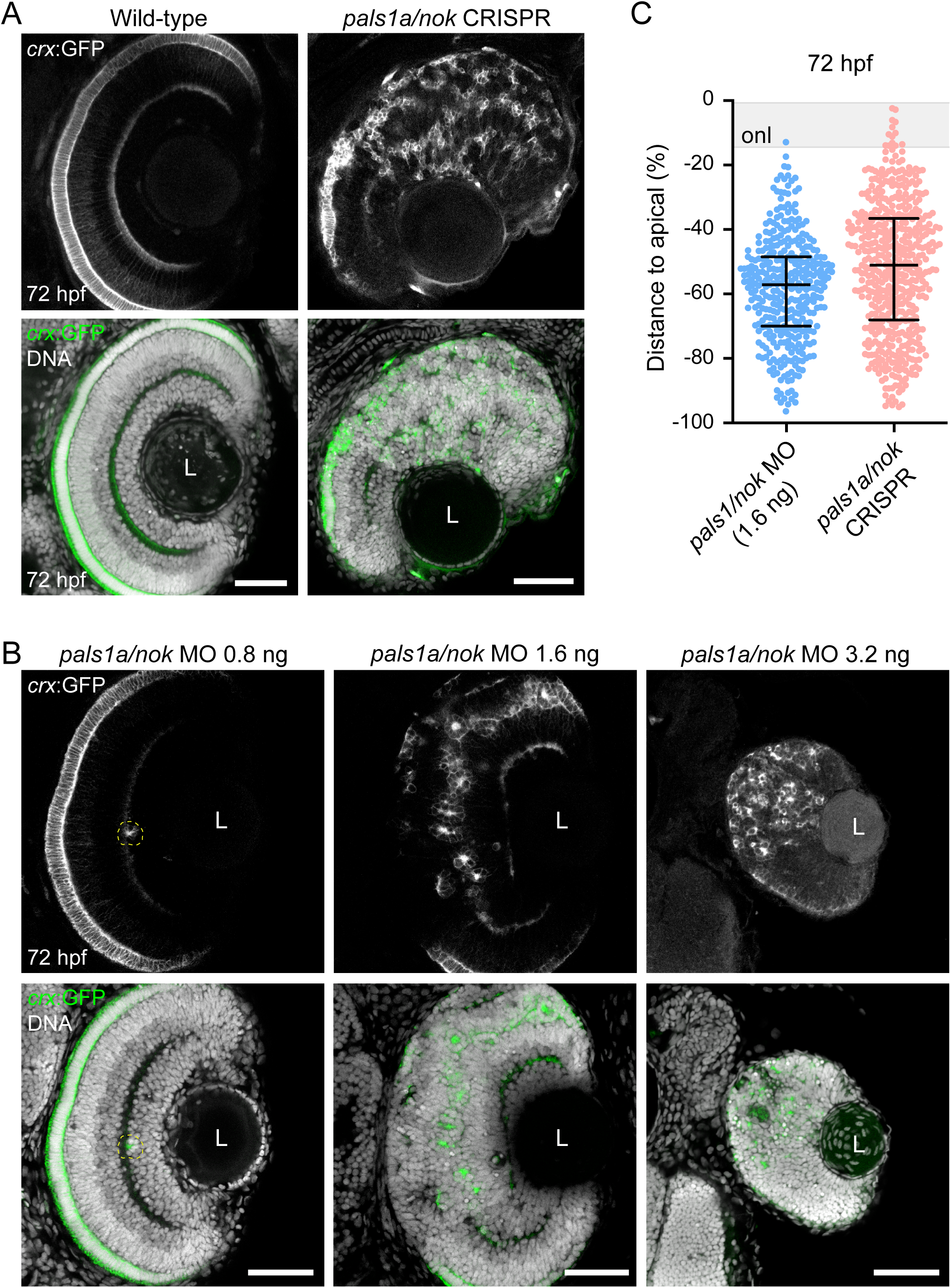
Photoreceptor positioning is disrupted upon *pals1a*/*nok* knockdown using morpholino oligomer injection and CRISPR/Cas9 F0 genome editing. **A.** Single confocal optical sections through the central retina of representative Tg(*crx*:GFP) embryos at 72 hpf. In the wild-type retina photoreceptors align as a continuous ONL along the apical surface, while in the *pals1a*/*nok* crispant *crx*:GFP-positive cells appear widely distributed across the retinal neuroepithelium, either as isolated cells or as elongated clusters, eventually subdivided into rosettes. **B.** Single confocal optical sections through the retinas of 72 hpf Tg(*crx*:GFP) embryos injected with three different doses of *pals1a*/*nok* MO. The lowest dose (0.8 ng) produces a mild phenotype, with only a few photoreceptors in displaced positions (dashed line). At 1.6 ng, embryos show a wide distribution of *crx*:GFP-positive cells across the retina, comparable to that observed in *pals1a*/*nok* crispants, and relatively low cell death. The highest dose (3.2 ng) results in delayed development and extensive cell death, particularly in the retina. **C.** Quantification of *crx*:GFP-positive cell positioning along the apico-basal retinal axis in *pals1a*/*nok* morphants (1.6 ng MO) and *pals1a*/*nok* crispants at 72 hpf. The distance from the cell nucleus to the apical surface (position 0) was normalized as a percentage of total retinal width. Each dot represents an individual cell. Approximately 50% of *crx*:GFP-positive cells localize between 48–70 % of the retinal width in morphants and between 40–68% in crispants, indicating broadly comparable distributions across both expression perturbation strategies. n: morphants, 359 cells from 13 embryos (4 independent experiments); crispants, 282 cells from 7 embryos (2 independent experiments). Median and interquartile range are shown. The gray band represents the approximate position of the outer nuclear layer (onl). L, lens. Scale bars: 50 μm.

As MOs offer the possibility of a wider dose-response range, we then injected embryos with different amounts of the *pals1a/nok* MO, seeking to find a situation in which the phenotype was adequate for following the differentiation of photoreceptor progenitors under hypomorphic conditions in which some cell adhesion could remain. When injecting 3.2 ng, we observed delayed development and extensive cell death, particularly in the retina (Fig. 1B). The lowest amount (0.8 ng), on the other hand, resulted in a very mild phenotype, in which only a few photoreceptors appeared in displaced positions. At 1.6 ng the retinal phenotype was relatively similar to that found in crispant embryos, with a wide distribution of photoreceptors across the retina, either accumulated in large groups or in smaller rosettes, and relatively low cell death (Fig. 1B). In these conditions, a quantification of *crx*-positive cells positioning in the apico-basal retinal axis showed broadly comparable distributions between morphants and crispants (Fig. 1C). Given the higher phenotypic consistency, the following experiments were all performed using 1.6 ng MO injection.

### *pals1a/nok* morphants display photoreceptor accumulations with preserved cell polarity and glial associations

To evaluate the effect of *pals1a*/*nok* deficiency on retinal organization, we performed MO-mediated gene knockdown in Tg(*crx*:GFP/*atoh7*:RFP) double transgenic embryos and analyzed retinal architecture at 48 and 72 hpf. Control embryos injected with standard MO showed properly positioned *crx*:GFP-positive photoreceptor progenitors at the outer nuclear layer (ONL), and a well-organized laminar structure was evident by 72 hpf, with clear segregation between *crx*:GFP-positive photoreceptors and *atoh7*:RFP-expressing RGCs (Fig. 2A). *pals1a*/*nok* MO-injected embryos showed early disruption of retinal architecture, with ectopic photoreceptor aggregates already visible at 48 hpf. Morphant retinas at 72 hpf showed prominent photoreceptor rosettes distributed across the retinal tissue with many RGCs mispositioned in apical or sub-apical regions. Nevertheless, the two cell populations remained segregated from each other (Fig. 2A). Higher magnification views of single confocal planes showed that *crx*:GFP-positive photoreceptor progenitors in morphant embryos formed ectopic rosettes in central retinal regions at both timepoints examined, rather than aligning along the apical surface as in controls (Fig. 2B). Cell height (apico-basal length) was significantly shorter in morphant photoreceptor progenitors than in controls at both stages: average 6.2 µm versus 8.5 µm at 48 hpf, and 7.5 µm versus 14 µm at 72 hpf (Fig. 2D).

**Figure 2.**
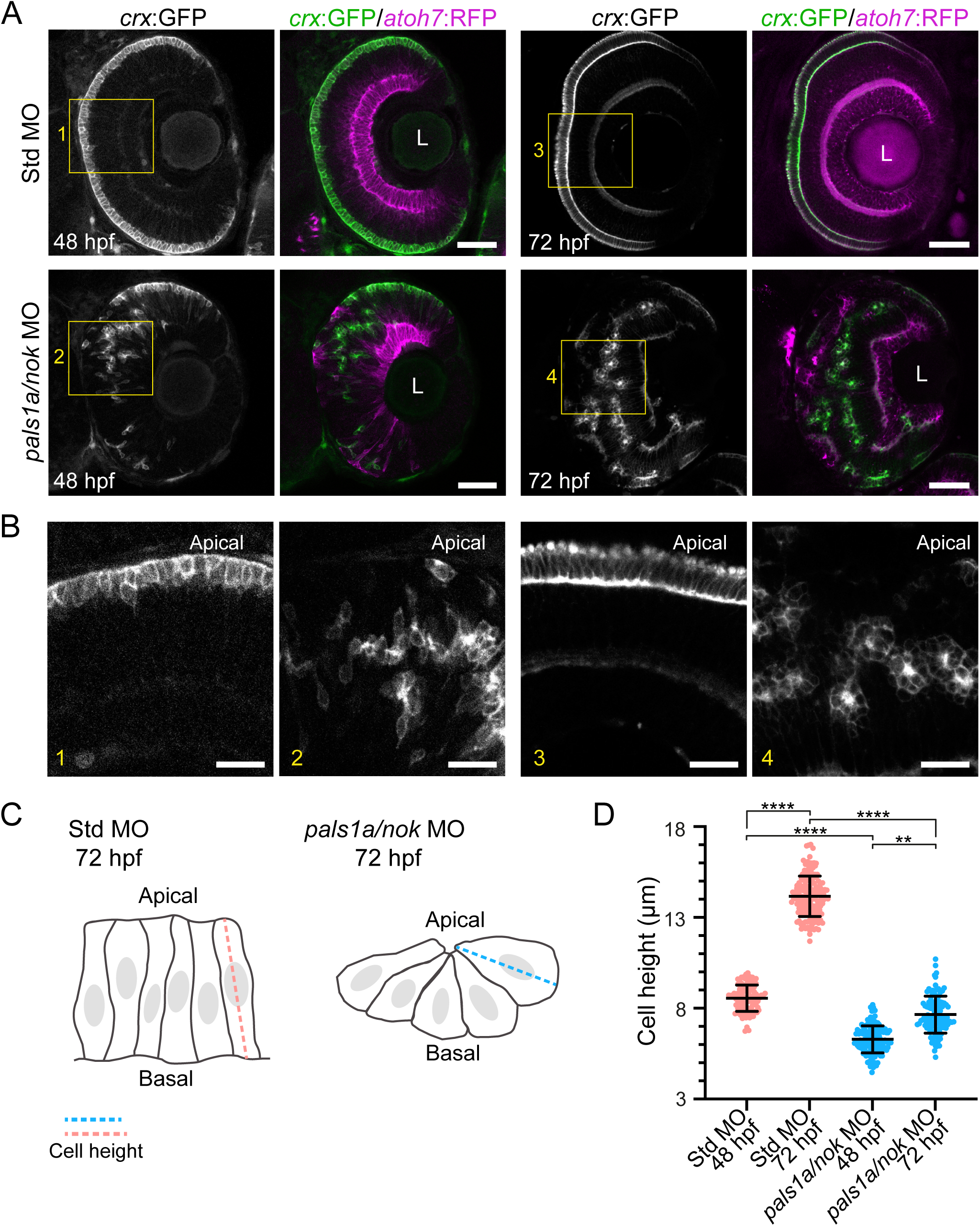
*pals1a*/*nok* morphants display photoreceptor rosettes with preserved cellular polarity. **A**. Retinas from standard MO-injected and *pals1a*/*nok* MO-injected Tg(*crx*:GFP/*atoh7*:RFP) double transgenic embryos. At 48 hpf, control retinas show emerging ONL organization with properly positioned *crx*:GFP-positive photoreceptor progenitors and *atoh7*:RFP-positive RGCs. Morphant retinas display early disruption with ectopic photoreceptor groups forming rosette-like structures. At 72 hpf, control retinas exhibit laminar organization with segregated cell types. Morphant retinas show architectural disruption with photoreceptor rosettes and mispositioned RGCs, but segregated cell populations. L, lens. **B**. Higher magnification views of the boxed regions from A. Apical orientation is indicated. Maximum intensity projection of 3 confocal planes (1 μm separation). **C**. Schematic of the cell height (apico-basal length) measurement, taken on single confocal planes as the distance between the apical and basal ends of each *crx*:GFP-positive cell. **D**. Comparative quantification of cell height (apico-basal length) of *crx*:GFP-positive cells. At 48 hpf, photoreceptor progenitors measure 8.5 μm on average in control embryos and 6.2 μm in morphants. At 72 hpf, photoreceptor progenitors in controls reach 14 μm, while in *pals1a*/*nok* morphants they reach only 7.5 μm. n: control 48 hpf, 107 cells from 5 embryos (2 independent experiments); control 72 hpf, 154 cells from 8 embryos (2 independent experiments); morphant 48 hpf, 132 cells from 5 embryos (2 independent experiments); morphant 72 hpf, 116 cells from 7 embryos (2 independent experiments). Median and interquartile range are shown; (**) p < 0.01, (****) p < 0.0001, Mann-Whitney test. Scale bars: A, 50 μm; B, 20 μm.

We asked ourselves whether the glial scaffold was also affected in morphant retinas and whether Müller glia maintained their associations with photoreceptors. Using the Tg(*gfap*:EGFP) transgenic line combined with immunolabeling for double cones (zpr-1 antibody) at 72 hpf, we observed in control retinas the characteristic radial morphology of Müller glia, with somata in the inner nuclear layer and processes extending to both the apical and basal retinal surfaces (Fig. 3A). At the apical border, Müller glia established tight associations with photoreceptors, forming the OLM. *pals1a/nok* morphant retinas showed tissue-level disorganization, with Müller glia somata ectopically positioned in both apical and basal regions (Fig. 3B). Despite this disruption, Müller glia extended processes that reached the central region of ectopic photoreceptor rosettes. OLM-like structures persisted at rosette centers, which indicates that the molecular interactions underlying glia-photoreceptor adhesion at the apical surface are preserved even without normal tissue lamination (Fig. 3B).

**Figure 3.**
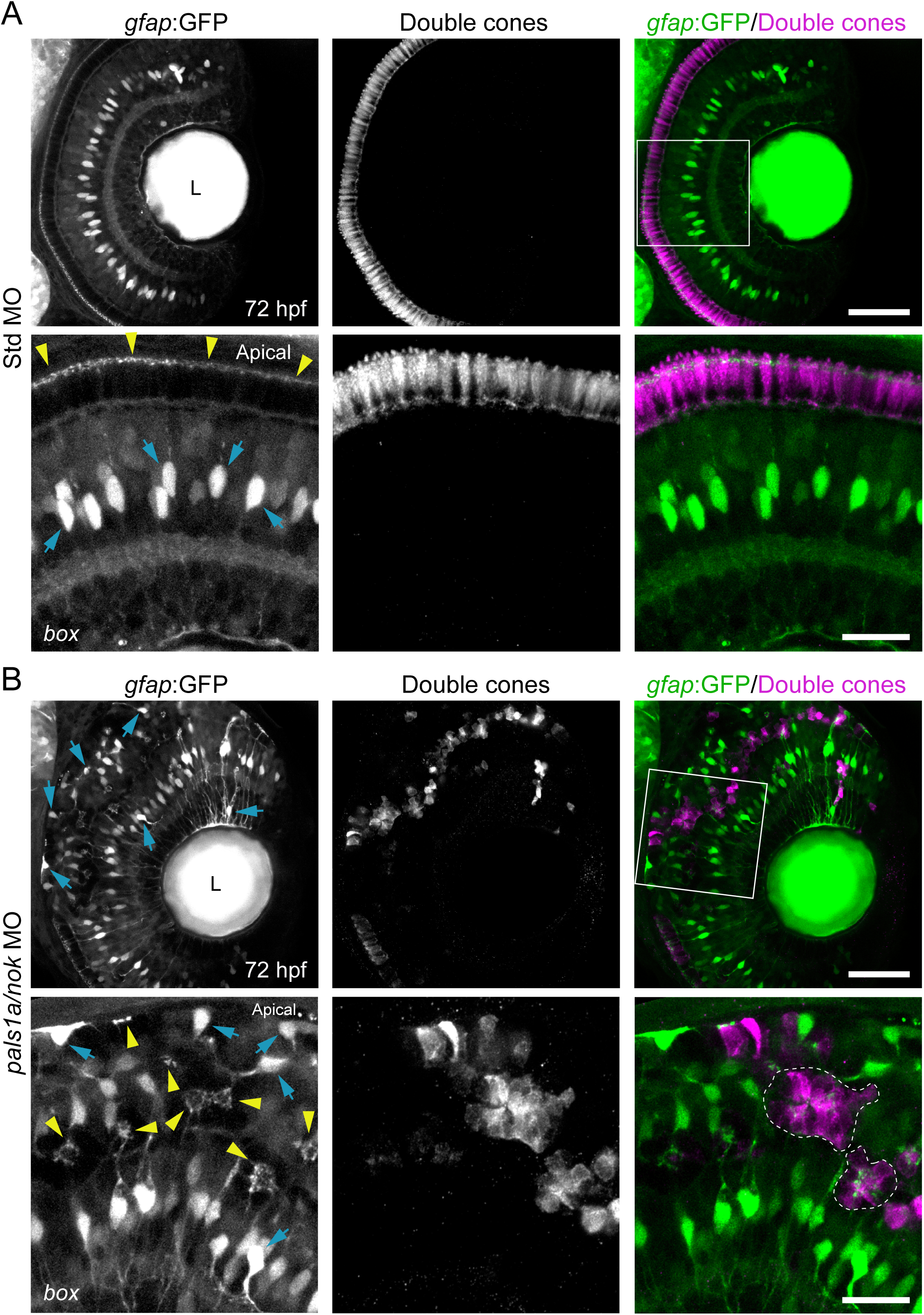
Müller glia maintain associations with photoreceptors in *pals1a*/*nok*-morphant retinas. **A.** Control retina showing normal Müller glia organization at 72 hpf. Transgenic *gfap*:GFP (Müller glia) embryos were immunolabeled with zpr-1 antibody (double cones). The lower image row shows a higher magnification of the boxed area, highlighting Müller glia somata (arrows) and apical OLM contacts (arrowheads). Müller glia exhibit a characteristic organization with apical and basal contacts; their somata localize to the inner nuclear layer. **B.** *pals1a/nok* morphant retina showing disrupted tissue organization. Low and high magnification images of the same optical sections show the ectopic positioning of Müller glia somata (arrows). Müller somata extend processes (arrowhead) toward the center of photoreceptor rosettes (dashed white outline), forming OLM-like structures. Maximum intensity projections of 5 confocal planes, 1 μm separation. L, lens. Apical orientation is indicated. Scale bars: overview panels, 50 μm; high magnification, 20 μm.

### Ectopic photoreceptor progenitors exhibit exploratory migration dynamics leading to rosette formation

To understand the cellular dynamics underlying rosette formation in *pals1a*/*nok* morphants, we performed time-lapse confocal imaging on Tg(*crx*:GFP/*atoh7*:RFP) double transgenic embryos starting at 36 hpf, a stage at which the first *crx*:GFP-positive photoreceptor progenitors become detectable. Low magnification views at the onset of imaging showed scattered *crx*:GFP-positive cells in the anterior region of morphant retinas, with the neural retina boundaries clearly identifiable by brightfield imaging (Fig. 4A). Time-lapse sequences documented how ectopic photoreceptor progenitors coalesced into rosettes (Fig. 4B). We tracked individual cells and observed that a *crx*:GFP-positive cell, initially adjacent to other *crx*:GFP-positive cells at the apical neuroepithelium at 36 hpf, migrated along the apical border. This cell underwent division at t = 160 min. By 300 min, two daughter cells were clearly visible and remained attached for approximately 50 min, after which both cells initiated divergent migratory trajectories toward *crx*:GFP-expressing regions: one daughter cell migrated along the apical retina toward a forming rosette, while the second moved through the tissue to incorporate into the same *crx*:GFP-positive cell group. Higher magnification imaging showed that during migration, cells established transient adhesive contacts with other *crx*:GFP-positive cells encountered along their path (Fig. 4C). We observed a tracked cell establishing transient adhesion with another *crx*:GFP-positive cell at the apical border and pausing migration for approximately 60 min during this adhesion period, while maintaining intense cortical activity. After separation, the cell completed its migration toward the rosette.

**Figure 4.**
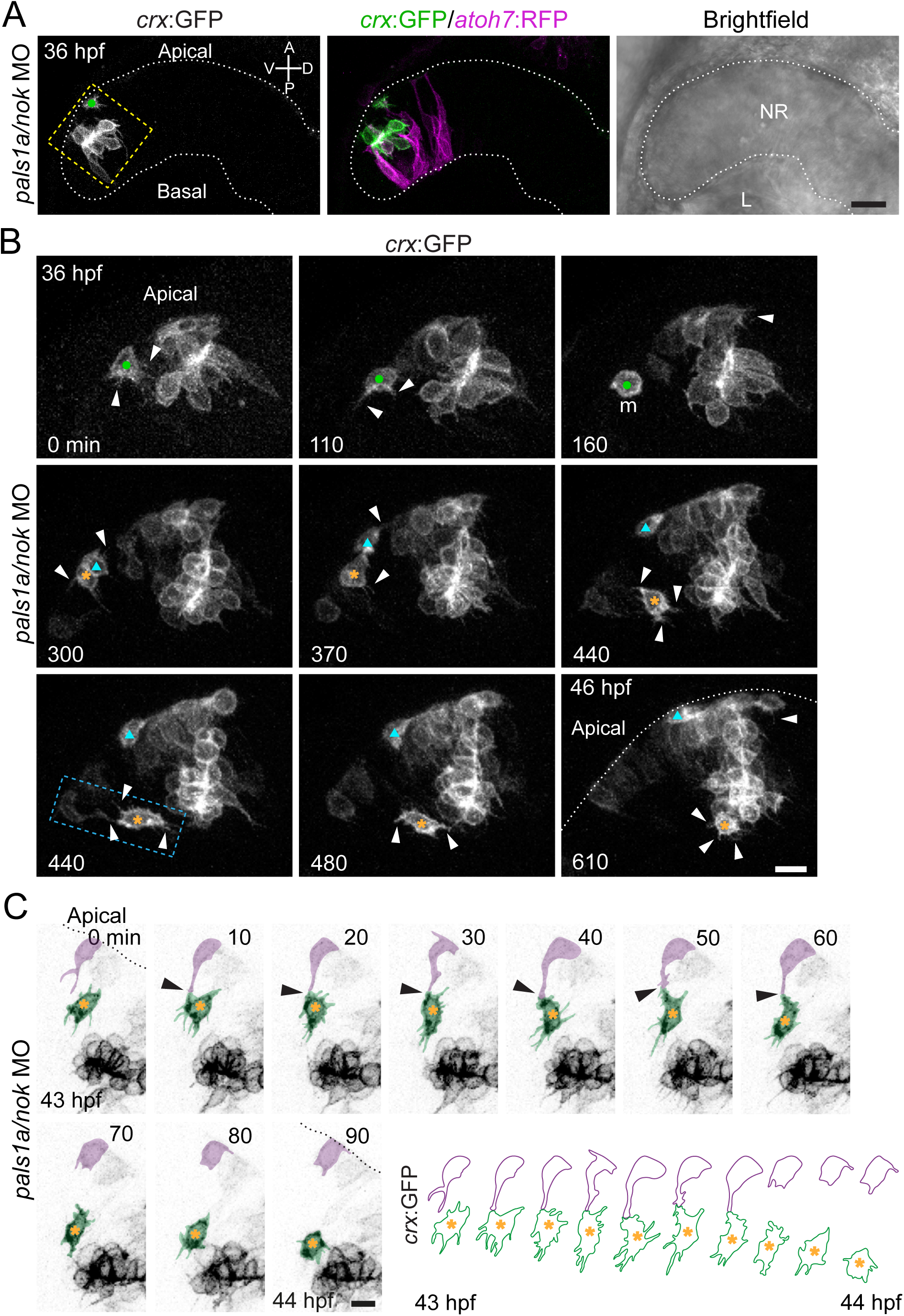
Effect of *pals1a*/*nok* knockdown on photoreceptor progenitor positioning dynamics. Confocal time-lapse imaging was performed on Tg(*crx*:GFP/*atoh7*:RFP) double transgenic embryos from 36 hpf. **A**. Low magnification confocal images at the beginning of the time-lapse, showing the anterior region of a *pals1a*/*nok* morphant retina. The dotted line marks the neural retina boundary. NR, neural retina; L, lens. Orientation axes are indicated (V, ventral; D, dorsal; A, anterior; P, posterior). **B**. Time-lapse sequence documenting rosette formation by ectopic photoreceptor progenitors (region delineated by the dashed box in A). A *crx*:GFP-positive cell (dot), initially located adjacent to other photoreceptor progenitors at the apical neuroepithelium at 36 hpf, moves along the apical border. At 160 min, the cell undergoes cell division (m). At 300 min, two daughter cells are clearly visible and remain contacting each other for approximately 50 min. Subsequently, both cells initiate divergent migratory trajectories toward *crx*:GFP-expressing regions: the cell with the triangle migrates apically, while the second cell (asterisk) moves to the center of the tissue; both cells ultimately incorporate into ectopic *crx*:GFP-positive cell groups. Arrowheads indicate cellular processes. Elapsed time is indicated in minutes. **C.** Time-lapse sequence showing higher magnification of a single cell from B (region delineated by the dashed box). The cell marked with an asterisk (and digitally highlighted) establishes transient adhesion with another *crx*:GFP-positive cell (digitally highlighted) located at the apical border. Arrowheads indicate the contact zone between both cells. The dotted line marks the apical retinal border. Although migration temporarily stops during this 60 min adhesion period, the asterisk-labeled cell maintains intense cortical activity. Subsequently, cells separate and the tracked cell completes its migration toward the rosette. Temporal resolution: 10 min. Digital representation below emphasizes the interaction between both cells and morphological changes, with cell trajectories and positions illustrated (colored lines and dots). Scale bars: A, 20 μm; B, 20 μm; C, 10 μm.

Focusing on the anterior-ventral retinal region, we performed quantitative analysis of the migratory behavior by tracking individual *crx*:GFP-positive cells from the onset of GFP detection (around 36-38 hpf) to their incorporation into rosettes (Fig. 5A). The displacement time was 356 ± 126 min (mean ± SD). Tracked trajectories showed that cells displayed apparently random movements with variable speeds and directions, although they did not translocate long distances overall. We compared accumulated distance (total path traveled) with Euclidean distance (effective displacement between starting and final positions) and found that the accumulated distance was considerably greater than the Euclidean distance in all cases, which is consistent with an apparently random movement pattern rather than directed migration (Fig. 5B). Instantaneous speeds averaged over the entire tracking period showed high inter-cell variability, which further supports the stochastic nature of this migratory dynamics (Fig. 5C).

**Figure 5.**
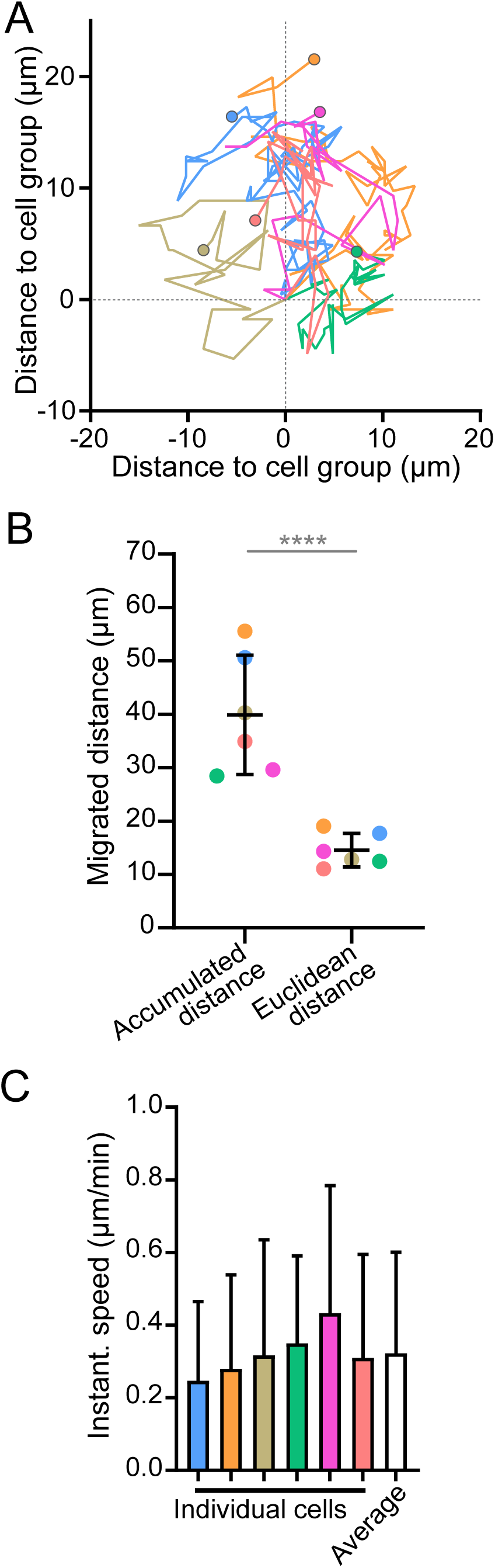
Quantitative analysis of exploratory migration dynamics in *pals1a*/*nok* morphants. Photoreceptor progenitor dynamics was quantified from time-lapse confocal observations. **A**. Tracked trajectories (x-y coordinates) of six *crx*:GFP-positive cells (3 embryos, 2 independent experiments) from the onset of GFP detection to incorporation into cell groups. All trajectories are aligned to the final position (0 μm in both axes); starting points are marked with colored dots. Cells displayed apparently random movements with variable speeds and directions despite not effectively translocating long distances. **B.** Comparison of total distance migrated (Accumulated distance) with effective displacement (Euclidean distance) in *pals1a*/*nok* morphants. Data correspond to cells analyzed in A, maintaining the color code. Mean ± SD; **** p < 0.0001. Accumulated and Euclidean distances (measured on the same cells) were compared with a paired Student’s *t*-test. **C.** Instantaneous speeds of photoreceptor progenitors from A (same color code), averaged over the entire tracking period. Each colored bar represents a single cell; white bar indicates the population average of instantaneous speeds.

### Retinal cells from *pals1a/nok* morphants maintain self-organization capacity with inverted topology in organoids

To test whether retinal cells from *pals1a/nok* morphants retain intrinsic self-organization capacity, we dissociated retinal cells from 24 hpf Tg(*crx*:GFP) embryos, reaggregated them in organoid culture and analyzed them after 48 hours. Control organoids show *crx*:GFP-positive photoreceptor progenitors preferentially positioned at the aggregate center (see Figs. 6A and 7A), as has been previously shown (Eldred et al., 2017). Immunolabelling for the apical polarity determinant aPKCζ indicates that in these retinal organoids, photoreceptor progenitors polarize into rosette-like structures, with aPKCζ accumulating at their innermost region (Fig. 6). F-actin accumulation around the central photoreceptor cell-cell contact regions also supports this idea (Fig. 7A). *pals1a/nok* morphant organoids showed a different spatial organization: photoreceptor progenitors redistributed toward peripheral regions of the aggregates, while F-actin localization at cell-cell contacts was maintained, this time at the organoid surface (Fig. 7A). Quantification of photoreceptor progenitor distribution confirmed this inversion, with the photoreceptor progenitor-to-center distance ratio shifted toward peripheral positioning in morphant organoids compared to relatively more central localization in controls (Fig. 7B). The spatial reorganization was not limited to photoreceptors. RGC distribution analyzed by zn-8 immunolabeling showed a complementary inversion: RGCs localized preferentially at the aggregate periphery in control organoids, but to central regions in *pals1a*/*nok* MO organoids (Fig. 7C). This preservation of the relative arrangement between cell types, despite the inverted radial topology, indicates that the mechanisms regulating relative cell-type positioning are maintained.

**Figure 6.**
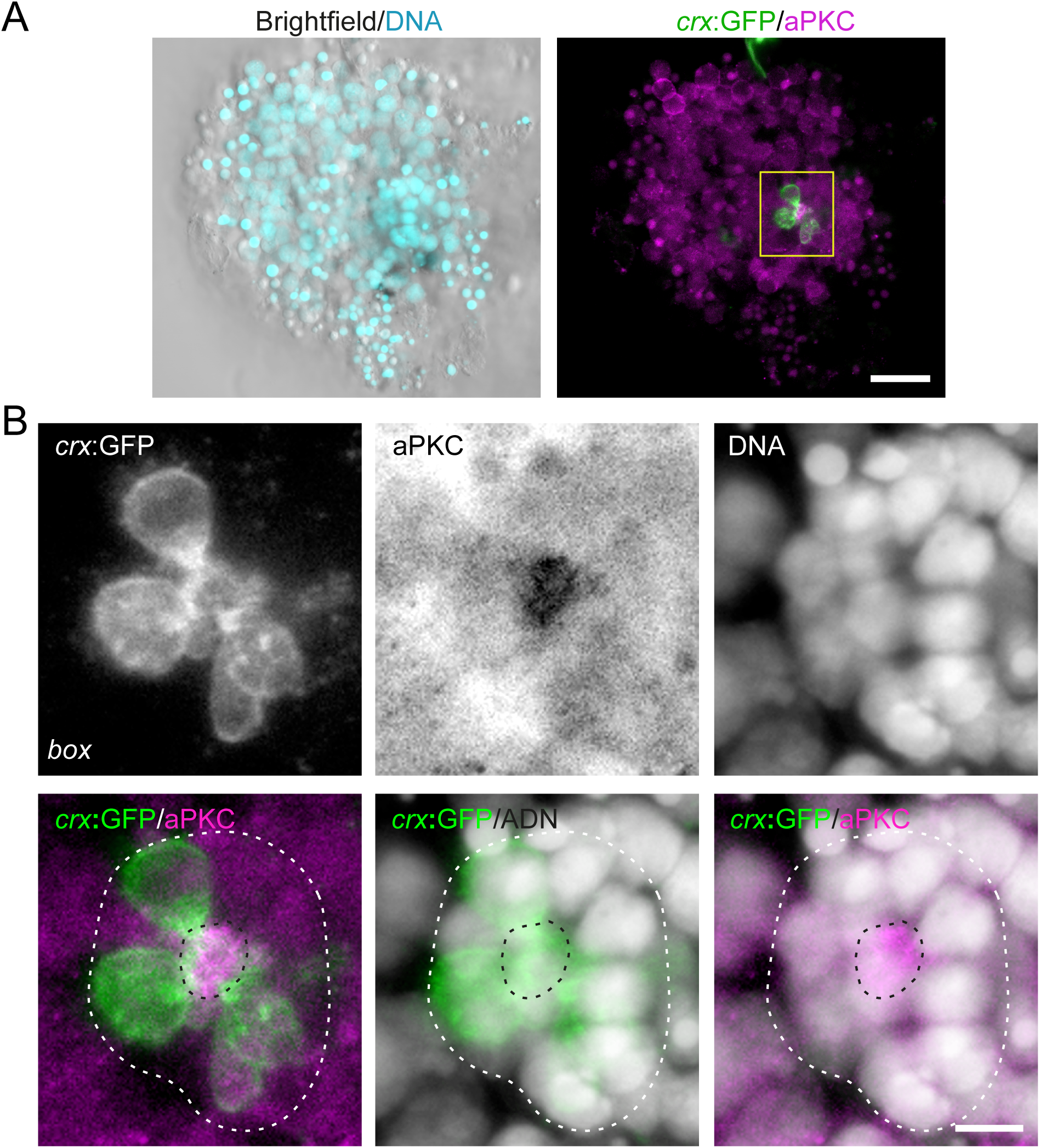
Apico-basal polarization of centrally-localized photoreceptor progenitors in retinal organoids. **A.** Retinal organoid generated from wild-type *crx*:GFP retinal cells and cultured for 48 h, immunolabeled for aPKCζ. The box outlines the region enlarged in B. **B.** Higher magnification views of the boxed region in A, showing a rosette-like structure. Black dotted lines indicate the apical border; white dotted lines delimit the rosette. Scale bars: A, 20 μm; B, 5 μm.

**Figure 7.**
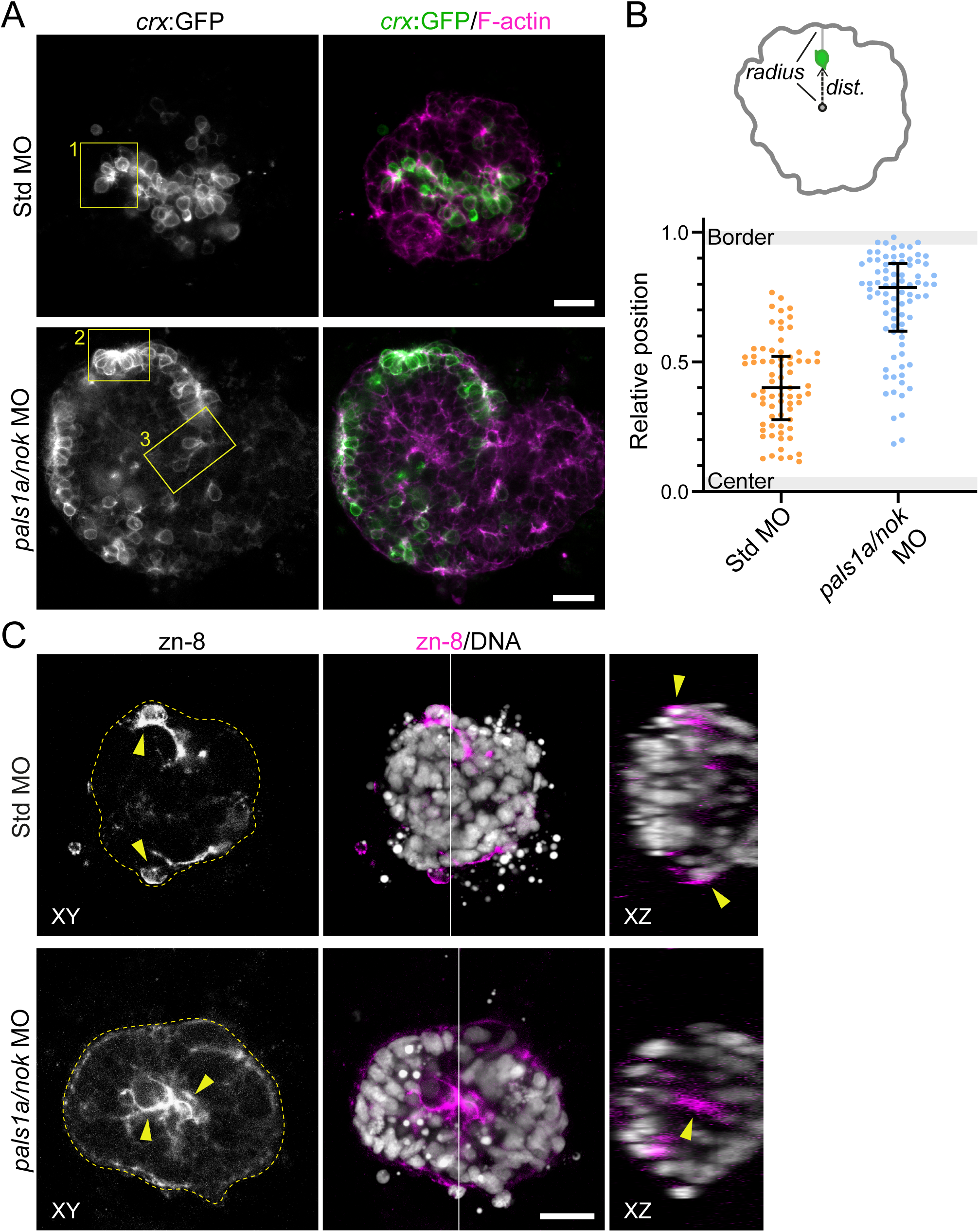
Retinal cells in *pals1a*/*nok*-morphant organoids maintain self-organization capacity but display inverted topology. **A.** Comparative analysis of retinal organoids generated from Tg(*crx*:GFP) embryos injected with standard or *pals1a/nok* morpholinos. Control organoids display compact architecture, with *crx*:GFP-positive photoreceptor progenitors positioned around the aggregate center. In *pals1a*/*nok* morphant organoids, photoreceptor progenitors (*crx*:GFP, green) redistribute towards peripheral regions. The yellow boxes indicate the regions shown at higher magnification in Figure 8. **B.** Quantification of photoreceptor progenitor distribution within organoids. The photoreceptor progenitor-to-center distance ratio (dist./radius) shows relative cell positioning, comparing peripheral (Border) versus central (Center) regions. Each dot represents an individual cell (n: 71 cells from 3 control organoids; n: 82 cells from 4 *pals1a*/*nok* organoids). Bars show median ± interquartile range. **C.** Spatial reorganization of RGCs: while in control conditions these cells tend to distribute at the organoid periphery, in *pals1a*/*nok* organoids they localize preferentially in central regions (yellow arrowheads). The white line represents the orthogonal view (XZ). Scale bars: 20 μm.

The analysis of both control and *pals1a/nok* morphant organoids showed that photoreceptor progenitors extended cellular processes (Fig. 8A). F-actin labelling revealed microfilament enrichment within cellular extensions in *pals1a/nok* morphant photoreceptor progenitors located at the cell aggregate periphery, where their separation from neighboring cells allowed unambiguous co-localization with TRITC-phalloidin labeling. In addition, in morphant organoids we could observe some individual *crx*:GFP-positive cells close to the periphery, which displayed an elongated cell process contacting an area bounded by microfilaments and devoid of nuclei (Fig. 8B). This morphology resembled that of photoreceptor progenitors attached to the retinal surface through an apical process during cell body translocation to the ONL (Aparicio et al., 2021).

**Figure 8.**
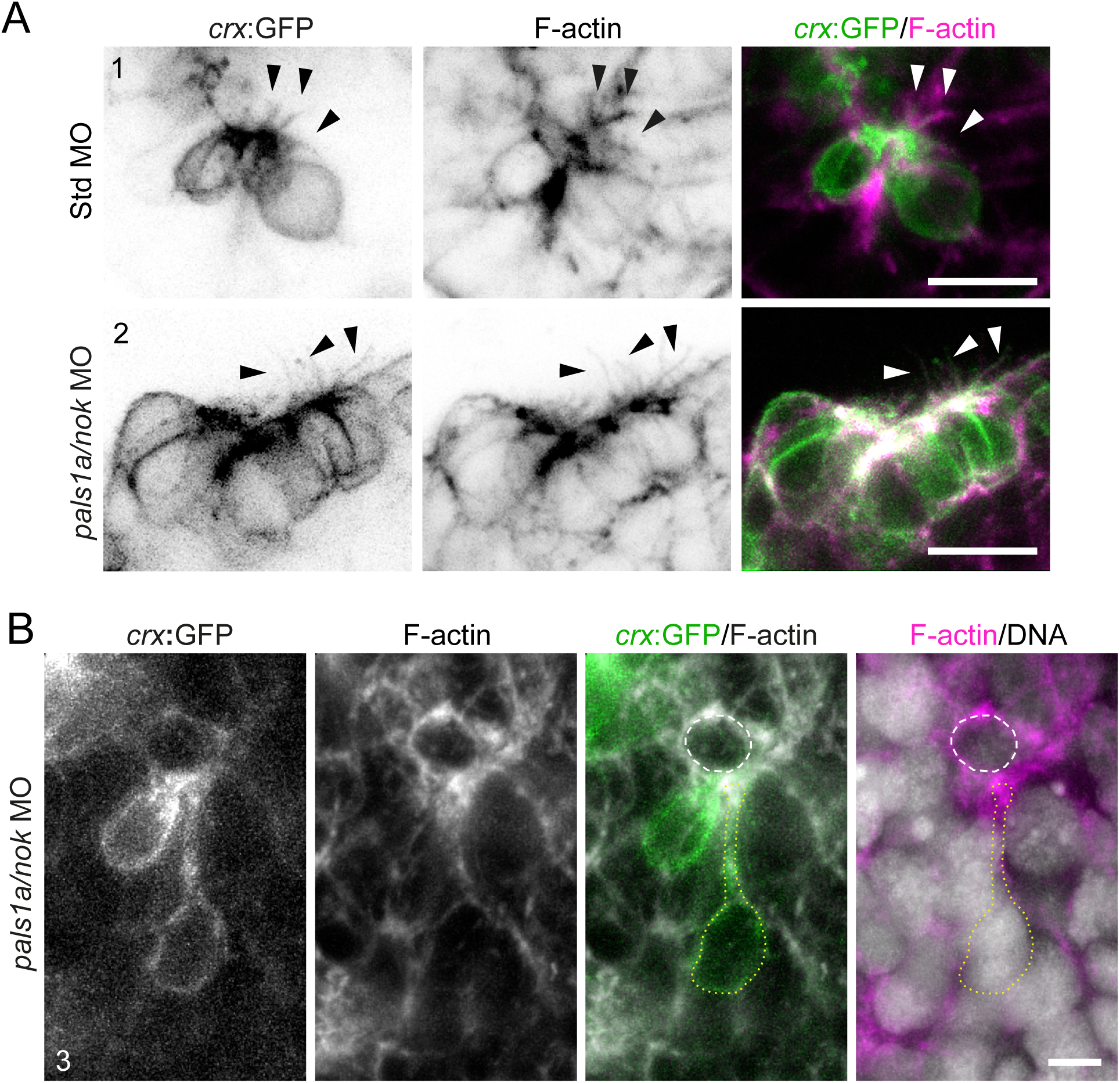
*pals1a/nok* morphant photoreceptor progenitors maintain early differentiation morphology and F-actin-rich apical cellular extensions in retinal organoids. **A.** Higher magnification views of the regions indicated by boxes 1 (control) and 2 (*pals1a/nok* morphant) in Figure 7A. The arrowheads indicate cellular extensions of *crx*:GFP-positive photoreceptor progenitors in both conditions. Maximum intensity projection of 4 confocal planes, 1 μm separation. **B**. Higher magnification view of the region indicated by box 3 (*pals1a/nok* morphant) in Figure 7A. The dashed circle indicates an “apical” surface lined by microfilaments and lacking nuclei. The dotted line marks a *crx*:GFP-positive cell with a cell process contacting this actin-rich surface. Maximum intensity projection of 5 confocal planes, 1 μm separation. Scale bars: A, 10 μm; B, 5 μm.

## Discussion

Following our previous observations of the behavior of *crx*-expressing photoreceptor progenitors in the Cadherin2-deficient zebrafish retina (Aparicio et al., 2021), we here aimed at better characterizing the importance of another polarized adhesion molecule, PALS1, regarding the dynamics of photoreceptor early differentiation and ONL formation. PALS1 is known to be central in the organization of retinal cells, and previous observations based on null mutant lines have shown a very dispersed distribution of photoreceptors across the retina (Wei et al., 2006; Wei and Malicki, 2002). In order to better adjust the severity of the defects, seeking to evaluate just the direct effect on the neural retina, we have used a validated morpholino oligomer (MO) knockdown protocol that allows for the obtention of a hypomorphic-like phenotype when compared to null mutant alleles (Zolessi et al., 2006).

*In vivo* confocal imaging of *pals1a*/*nok* morphants showed a very similar dynamics to what we described for the *cdh2*-deficient embryos. Although some photoreceptors organized into short stretches of apically localized ONL, most of the initially dispersed, unpolarized *crx*-positive cells exhibited exploratory migratory behaviors, to eventually coalesce into ectopic cell groups localized inside the retina. These groups appeared large and elongated initially, to eventually fragment into smaller round groups with the apical border of differentiating photoreceptors towards the inside (“rosettes”), as reported for the *cdh2* morphants (Aparicio et al., 2021). Quantitative tracking of these trajectories between 36 and 46 hpf yielded instantaneous speeds of around 20 µm/h (0.32 µm/min) for *crx*-positive cone progenitors, higher than the ∼5 µm/h reported for rod precursors at ∼50 hpf (Wei et al., 2006); this difference likely reflects the earlier developmental window, the retinal region sampled and the higher temporal resolution of our imaging.

The main known interactions and functions of PALS1 are related to its integration of the Crumbs complex. Three of the five members of the adhesion protein family Crumbs (Crb) found in zebrafish have been shown to be important for photoreceptor differentiation and maintenance: Crb1, Crb2a and Crb2b (Omori and Malicki, 2006). While the others are more related to later, post-mitotic stages of photoreceptor differentiation, Crb2a appears more relevant at early stages, its mutant *ome* having an early retinal polarity phenotype very similar to that of *pals1a*/*nok* (Malicki and Driever, 1999). Nevertheless, the similarities between phenotypes of *pals1a*/*nok* and *cdh2* are not surprising, since both have functions in apical cell adhesion, and both have been previously shown to modulate cell proliferation in the retina (Yamaguchi et al., 2010).

Work from the Zou lab has shown that there is actually a functional interaction between the two adhesion complexes in zebrafish embryo retina and other epithelia. First, Crb2a is essential for the establishment of an apical adherens junction, and is also necessary for the proper localization of PALS1 at the apical border, from 48 hpf (Lin et al., 2020). Second, PALS1 interaction with the small GTPase Rab11 was shown to be necessary for Cadherin2 trafficking to the apical plasma membrane in lens epithelial cells, hence directly collaborating in the formation of the zonula adherens (Hao et al., 2020). These observations point to a central role of PALS1 in establishing apical cell adhesions in the zebrafish retina, and position its action upstream to that of Cadherin2. Interestingly, the rosettes formed in *cdh2* mutants appear to lack Müller glia apical processes (Wei et al., 2006), which was expected given that the zonula adherens configuring the OLM is established between glial and photoreceptor cells. Our present results, however, show that the internal cell groups and rosettes in the hypomorphic *pals1a*/*nok* morphants have a very highly organized mesh-like structure formed by Müller glia processes at their cores. Altogether, these observations indicate essential differences between the two adhesion complexes. This could be due to the observed restriction of PALS1 to photoreceptors (Wei et al., 2006) or because of its role in maintaining the integrity of the RPE, which in turn was shown to be essential for neural retina organization (Zou et al., 2008).

Retinal cell aggregates and organoids have traditionally been another way of studying the mechanisms of retinal layer formation. To further characterize the cellular basis of the *pals1a*/*nok* morphants phenotype, we isolated retinal progenitors to generate organoid cultures, obtaining the expected spontaneous layered organization with photoreceptors inside and RGCs around the surface (Eldred et al., 2017). Previous studies using chick embryo retinal cells have shown a similar order of organization, except that photoreceptors were grouped in multiple internal rosettes (Layer and Willbold, 1994; Moscona, 1961). Our present observations indicate that the internal photoreceptor accumulation in zebrafish retinal organoids is also a rosette-like structure, with apical cell borders towards the inside. The usual presence of only one rosette in this case could be related to a smaller cell aggregate size in our experimental setup.

This organization, with photoreceptors inside and RGCs outside, can be explained by the differential adhesion hypothesis, which implies that if there are no other signals involved, different cell types aggregated in culture tend to sort out with more adhesive cells in the inside (Steinberg, 1970). We show here that PALS1-deficient retinal cells retained intrinsic self-organization capacity in organoids, albeit with an inverted topology when compared to the control situation (i.e., with photoreceptors on the surface). Mechanistically, this inversion is consistent with the same differential-adhesion logic: because PALS1 promotes apical adhesion and the apical trafficking of Cadherin2 (Hao et al., 2020), its reduction would lower the relative adhesiveness of photoreceptor progenitors and displace them toward the aggregate surface. Photoreceptors differentiating in organoids extended F-actin-enriched processes resembling the apically localized tangential processes described *in vivo* (Aparicio et al., 2021; Hu and Masai, 2025; Sharkova et al., 2024). Thus, interestingly, in the PALS1-deficient retinal organoids, photoreceptors appear to be able to regain the capacity to organize into an ONL-like structure.

Albeit inverted in orientation, the maintenance of photoreceptor polarization in *pals1a*/*nok* morphant organoids supports the idea that these cells retain fundamental aspects of their differentiation and polarization program, including cell cortex and cytoskeletal organization associated with early morphogenesis, independently of normal tissue architecture. Interestingly, a similar layer order and polarity reversion was obtained in chick retinal aggregates when co-cultured with a monolayer of RPE cells separated by a 0.4 μm filter (Rothermel et al., 1997). Since a great part of the neural retina phenotype of PALS1 deficiency in whole embryos is related to the disruption of the RPE (Zou et al., 2008), the apparent “normalization” of neuronal localization in our *pals1a/nok* morphant aggregates could be due to an effect on these cells. However, not only RPE cells (which localize at the center of the organoid) are usually very few in these culture conditions, but it was also demonstrated that their absence does not affect layer organization (Eldred et al., 2017). Hence, the layer re-organization in these organoids is most probably a direct consequence of reducing PALS1 in neural retina cells.

Altogether, our results show an essential role for PALS1 in the regulation of photoreceptor positioning and ONL formation in the zebrafish retina, independently of the mechanisms ensuring cell-type laminar segregation and cell polarization. They also reinforce the idea that, despite phenotypic similarities, there are functional differences in the cell adhesion mediated by PALS1-Crumbs and Cadherin2 in the developing neural retina. Even when PALS1 is strongly reduced, photoreceptor progenitors retain their identity, polarize, establish OLM-like apical contacts with Müller glia and self-organize into cell groups; what fails is the apical positioning and stable anchoring of these cells. Since PALS1 is part of the apical adhesion machinery, its depletion would lower the relative apical adhesiveness of photoreceptor progenitors, offering a plausible explanation, in terms of differential adhesion, for their relocation from the apical surface to the retinal center in intact embryos, and from the center to the surface in organoids. PALS1 thus appears to set the position that photoreceptors acquire within their tissue environment, acting at least in part through apical adhesion, rather than being strictly required for an ordered, polarized architecture to emerge.

## Acknowledgements

We thank Kristen Kwan and Rachel O.L. Wong for sharing plasmids and/or fish lines. We also thank Camila Davison for her guidance in establishing the retinal organoid cultures, and Gisell González for her invaluable and continuous support with fish care. The authors gratefully acknowledge the Advanced Bioimaging Unit at the Institut Pasteur Montevideo for their support and assistance in the present work. This work was partly funded by ANII-FCE, Uruguay, grants to FRZ (1_1_2014_1_4982; FCE_1_2021_1_166427); SNB PhD fellowship to GA; Programa de Desarrollo de las Ciencias Básicas (PEDECIBA, Uruguay).

## Declarations of interest

The authors declare no conflicts of interest.

## Author contributions

GA: Conceptualization; Data curation; Formal analysis; Investigation; Methodology; Resources; Validation; Visualization; Writing - original draft; Writing - review & editing. FRZ: Conceptualization; Data curation; Formal analysis; Funding acquisition; Investigation; Project administration; Resources; Supervision; Validation; Visualization; Writing - review & editing.

## Notes

### Competing Interest Statement

The authors have declared no competing interest.

