## Supplementary for "Reduction of PALS1/Nok disrupts retinal lamination through altered cell positioning while preserving photoreceptor self-organization capacity"

**Supplementary material**

**Supplementary Table 1.** Antisense morpholino oligonucleotides used in this study.

| Morpholino | Target gene | Sequence (5'→3') | Reference |
|---|---|---|---|
| *pals1/nok* | *pals1a/nok* | CTTTCTGCATGGTGTGAGGGCTGAG | Zolessi et al., 2006 |
| Standard control | — | CCTCTTACCTCAGTTACAATTTATA | Gene Tools; Moulton, 2017 |
| p53 | *tp53* | GCGCCATTGCTTTGCAAGAATTG | Robu et al., 2007 |

Antisense morpholino oligonucleotides (Gene Tools, LLC) used in this study. The *pals1a/nok* and p53 morpholinos are translation-blocking oligonucleotides complementary to the region spanning the translation initiation site of their respective target transcripts (*pals1a/nok* and *p53*, respectively). The *pals1a/nok* morpholino is a previously validated reagent that produces a phenotype milder than the null mutant (Zolessi et al., 2006). The p53 morpholino was co-injected to suppress non-specific, p53-mediated apoptosis associated with morpholino microinjection (Robu et al., 2007). A standard control morpholino with no known target in the zebrafish genome was used as a negative control (Moulton, 2017). Morpholinos were resuspended in nuclease-free water and microinjected into the yolk of 1- to 4-cell stage embryos.

**Supplementary Table 2.** gRNAs used for CRISPR/Cas9 F0 mutagenesis of mpp5a gene (*pals1/nok)*.

| gRNA | Guide sequence (5'→3') | Exon | Genomic position (chr17) | Strand | Score | Encoded protein region |
|---|---|---|---|---|---|---|
| gRNA 1 | GGGGCTCTGAAGACCACCTG | 3 | 34.268.412–34.268.434 | – | 83 | Upstream of L27 domain |
| gRNA 2 | GGTCCGTCAGGGGGTAGGG G | 4 | 34.272.016–34.272.039 | – | 71 | L27 domain |
| gRNA 3 | GGCGAGACTCTCACCCAGTG | 6 | 34.279.851–34.279.873 | + | 70 | L27–PDZ linker region |
| gRNA 4 | GGCAGTCTCTCCTCCCCACT | 6 | 34.279.864–34.279.886 | – | 64 | L27–PDZ linker region |

Guide sequences (20-nt DNA-binding sequence of each gRNA, complementary to the genomic target; scaffold adaptors not shown) of the four gRNAs designed against *pals1a/nok* (NCBI Reference Sequence NM_194363.1; Ensembl gene ENSDARG00000006272) using CRISPRscan (Moreno-Mateos et al., 2015). Genomic sequence corresponds to chromosome 17 of the GRCz10/danRer10 zebrafish assembly. The targeted exons (3, 4 and 6) are shared by the three annotated splice variants of the locus. The four gRNAs span contiguous regions encoding the N-terminal portion of *pals1a/nok*, encompassing the L27 domain and the linker region between the L27 and PDZ domains. *Score*, CRISPRscan predicted cleavage activity score. Off-target prediction returned 0 sites in the seed region and 0 total off-target sites for all four gRNAs. The four gRNAs were pooled in equal volumes (500 ng/µL total; 125 ng/µL each) and assembled into ribonucleoprotein complexes by mixing 1:1 (v/v) with HiFi Cas9 Nuclease V3 (IDT, 1081061; 10 µM), yielding a final injection solution of 5 µM Cas9 and 250 ng/µL total sgRNA (62.5 ng/µL each). 1.5 nL of this complex was injected into the yolk of 1- to 4-cell stage embryos, following Wu et al. (2018).

Each guide sequence was incorporated into a target-specific forward primer carrying the T7 promoter at its 5' end (5'-taatacgactcactata-*[guide sequence]*-gttttagagctagaa-3') and a 3' tail complementary to a generic reverse primer that provides the constant tracrRNA-like scaffold of the sgRNA (Moreno-Mateos et al., 2015; Varshney et al., 2015). Generic primer sequence:

5'-AAAAGCACCGACTCGGTGCCACTTTTTCAAGTTGATAACGGACTAGCCTTATTTTAACTTGCTAT-3'

**Supplementary Videos**

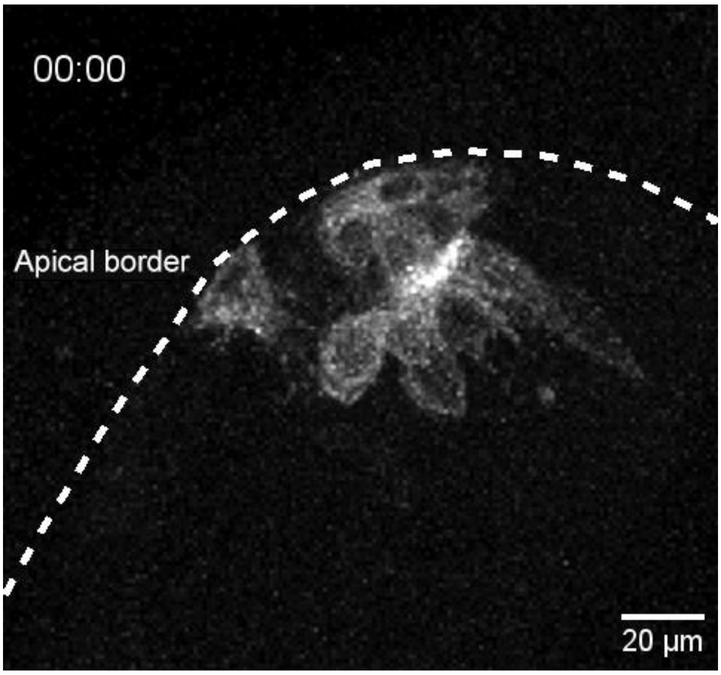


**Supplementary Video 1 (related to Figure 4B).** Confocal time-lapse sequence showing the process of ectopic *crx*:GFP-positive cell group formation by photoreceptor progenitors in *pals1a/nok* morphant retinas. Color dots indicate the migration of a progenitor that eventually divides; both daughter cells incorporate into a cell group. Temporal resolution: 10 min. See Fig. 4B for selected time points.

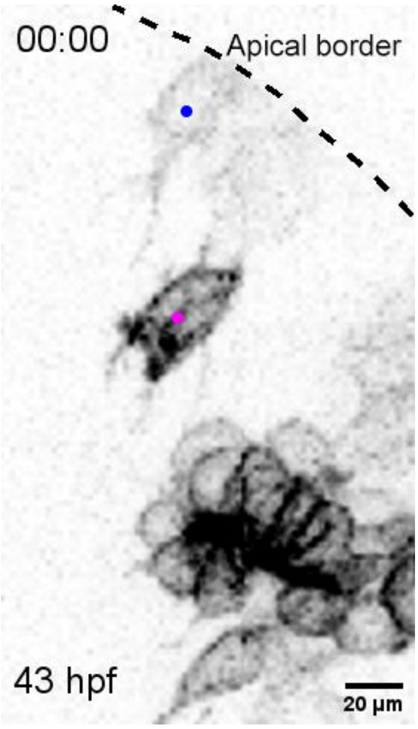


**Supplementary Video 2 (related to Figure 4C).** Time-lapse sequence from the same experiment shown in Fig. 4B, illustrating the dynamic interaction between photoreceptor progenitors in *pals1a/nok* morphant retinas. The cell marked with a magenta dot establishes a transient adhesion with another *crx*:GFP-positive cell (blue dot) located at the apical border of the retina. Arrowheads indicate the contact zone between both cells. Temporal resolution: 10 min. See Fig. 4C for selected time points.

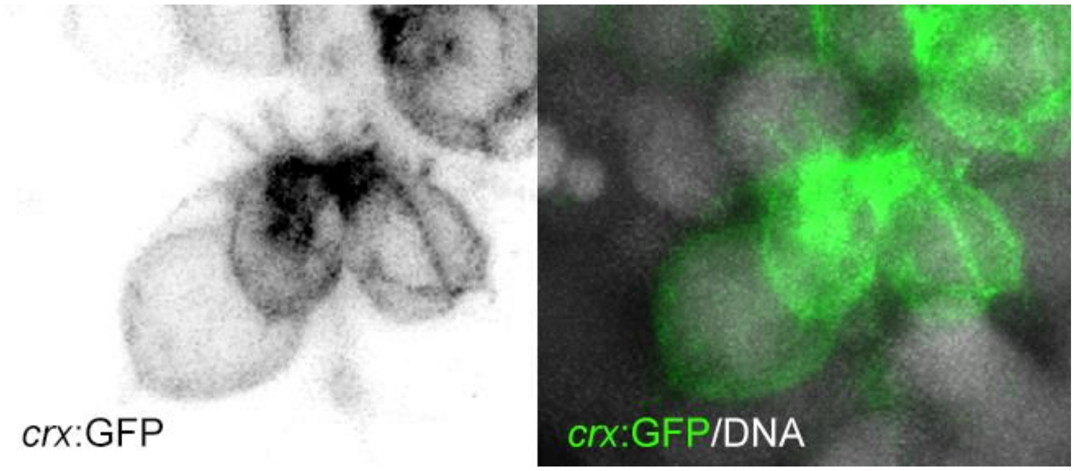


**Supplementary Video 3 (related to Figure 7A Std MO).** Three-dimensional reconstruction of *crx*:GFP-positive cells displaying polarized cellular extensions in retinal organoids derived from Standard MO-injected embryos. See Fig. 7A for further reference.

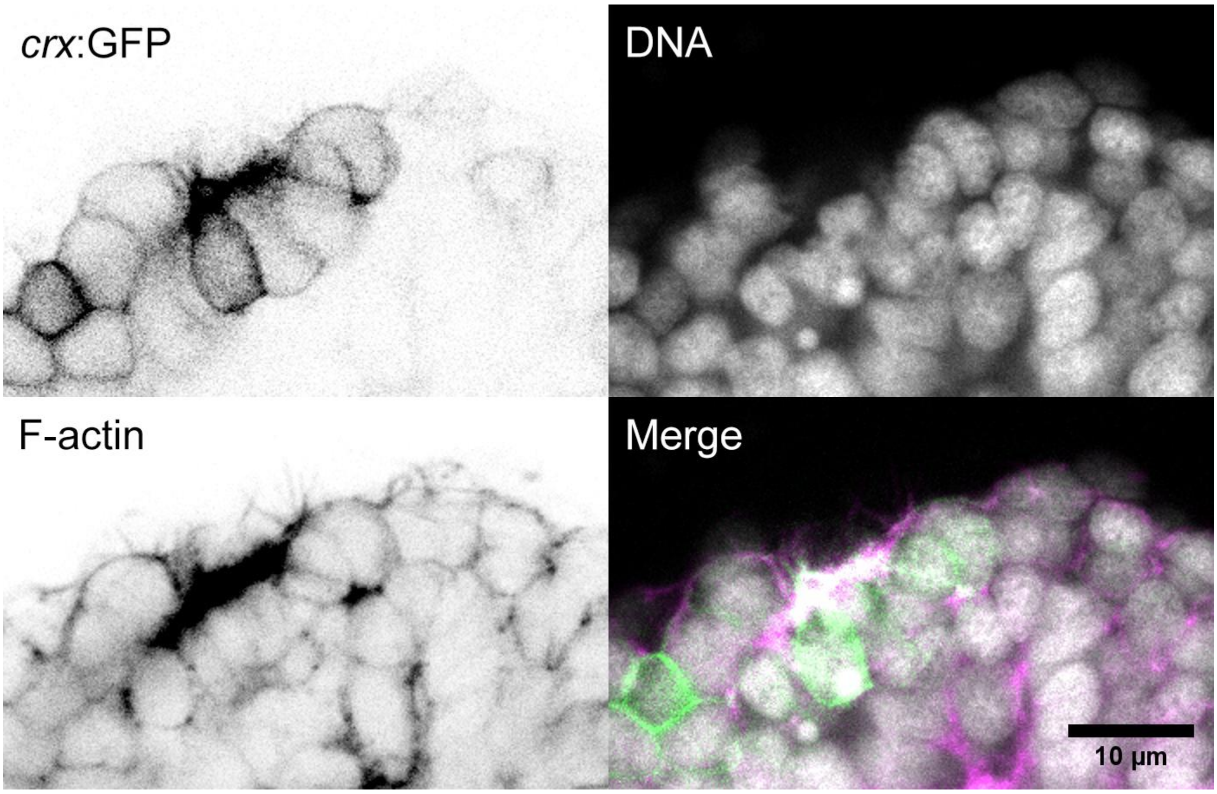


**Supplementary Video 4 (related to Figure 7A *pals1a/nok* MO).** Animation of serial confocal optical sections (z-planes) of *crx*:GFP-positive cells in retinal organoids derived from *pals1a/nok* morphant embryos, labeled with TRITC-phalloidin (F-actin) and methyl green (DNA). See Fig. 7A for further reference.
